# Actin nucleation promoting factors drive Arp2/3 dependent endosomal microautophagy

**DOI:** 10.64898/2026.07.09.737473

**Authors:** Satya Surabhi, Andreas Jenny

## Abstract

Autophagy is a catabolic process that degrades damaged organelles and aggregation-prone proteins and plays key roles during development and in maintaining cellular homeostasis. It can be induced by stress including starvation, oxidative stress, or accumulation of misfolded proteins. Autophagy declines with age and there is great interest in manipulating autophagy to improve neurodegenerative diseases, as its stimulation shows promise to improve diseases including Huntington, Alzheimer, and Parkinson. Endosomal microautophagy (e-MI) is a type of autophagy in which cytosolic proteins are delivered to late endosomes and degraded upon incorporation into intraluminal vesicles of multivesicular bodies. Here, we report that the actin nucleation-promoting factors (NPFs) known to activate the Arp2/3 complex to promote branched actin assembly can alter the dynamics of e-MI. We found that upon stress exposure, overexpression of the NPFs WASp, Wash, or SCAR results in an expedited induction of e-MI. Strikingly, Wash is uniquely required for physiological e-MI induction implying that NPFs are not functionally redundant for e-MI. We show that the WASH complex regulates e-MI on late endosomes acting via Arp2/3 and thus likely branched actin. Surprisingly, the regulation of e-MI by Wash is independent of retromer that is known to recruit Wash to early endosomes for its role in recycling of membrane proteins and rather reflects a novel degradative aspect of Wash function. Taken together, we identified a novel function of NPFs as upstream regulators of e-MI that could be used to activate e-MI ectopically to improve aggregate clearance during neurodegeneration.

## Introduction

Diseases such as cancer, cardiovascular diseases, and dementias are strongly affected by a decline in proteostasis. Autophagy is a major catabolic process that degrades damaged organelles and long-lived, aggregate-prone proteins in lysosomes and is critical to prevent neurodegenerative disorders, ^1–3^. For example in diseases, such as Alzheimer’s, Parkinson’s, amyotrophic lateral sclerosis (ALS), and polyglutamine (polyQ) diseases, protein aggregates in the brains of patients have long been recognized as a common pathological feature and promotion of autophagy to counteract aggregates is viewed as a potential therapy ^4–6^. To date, three main types of autophagy have been described. During macroautophagy (MA), cytosolic components are engulfed in autophagosomes that fuse with lysosomes for degradation ^7–9^. Chaperone-mediated autophagy (CMA) targets soluble cytosolic proteins with a KFERQ-like motif that is recognized by the heat shock cognate protein 70 (Hsc70, aka. HSPA8) and is required for lysosomal translocation by LAMP2A (Lysosome associated membrane protein 2A) ^10–14^. Both, MA and CMA can be strongly induced by cellular stress, often in a temporally distinct manner with CMA following MA. During endosomal microautophagy (e-MI), cytosolic proteins are targeted either in bulk or in a KFERQ-motif dependent manner to late endosomes (LE), where they are internalized into intraluminal vesicles of multivesicular bodies (MVBs) for degradation in LEs or lysosomes ^15–17^. Importantly, various forms of e-MI all require several components of the ESCRT (endosomal sorting complex required for transport) machinery. ESCRT mediates the invagination events during MVB formation and likely relies on specialized mechanisms of cargo organization and membrane deformation at endosomal surfaces that remain unresolved ^16^.

Much remains to be learned about the regulation and physiological role of e-MI (reviewed in ^16^). While e-MI contributes to protein quality control and can process aggregate-prone proteins in mammals, it mostly is thought to be a constitutive process that is even repressed upon stress such as prolonged starvation ^18–20^. An age-associated decline in e-MI in mice is accompanied by a compensatory increase in protein secretion through exosomal vesicles, suggesting that cargo normally cleared by e-MI is instead redirected into the extracellular space, which raises the possibility that restoring e-MI activity could represent a therapeutic strategy for neurodegenerative disorders ^21^. Studies in *Drosophila* have demonstrated that e-MI can occur in a constitutive manner but also be stress induced^22, 23^. Constitutive e-MI is involved in maintaining synaptic fitness via control of the turnover of synaptic proteins ^23^. In flies, e-MI is best characterized in the larval fat body that has functions akin to mammalian liver and adipose tissue, where it is induced by selective forms of stress including starvation and ROS. It requires the ESCRT machinery and, at least in part, Hsc70-4 ^22, 24^. e-MI in the FB is genetically and temporally distinct from MA. It’s induction requires prolonged (>12h) starvation relative to MA that peaks after 4h ^22, 25, 26^ and does not require the core MA genes *Atg5*, A*tg7*, and *Atg12* ^22^. Although the cause for the distinct activation times is unknown, starvation induced e-MI and MA are both coupled to inactivation of TOR kinase (target of Rapamycin), a conserved nutrient-sensing kinase that regulates cellular growth and autophagy ^22, 27^.

Here we report novel function for actin nucleation promoting factors (NPFs) regulating the timing of e-MI in fat body of *Drosophila* through Arp2/3 mediated actin branching. The Arp2/3 complex is a seven-subunit protein complex that binds to the side of a pre-existing actin filament and nucleates a new “daughter” filament at a characteristic 70° angle, thereby generating branched actin networks that drive membrane protrusion, vesicle formation, and intracellular transport ^28, 29^;Khaitlina, 2014 #567}. Because this process is inefficient, Arp2/3 activity depends on a family of NPFs which stimulate actin polymerization to physiologically relevant levels ^30, 31^. The NPF protein family includes WASp, WASH, WAVE/SCAR, and WHAMY, a WASp-family paralog. WASp, SCAR, and WASH contain a conserved VCA (verprolin-homology/central/acidic) domain that simultaneously recruits actin monomers and activates the Arp2/3 complex, coupling upstream signaling to localized actin assembly (Fig. S1) ^31–34^. The WASp related subfamily of WHAM proteins lacks a VCA domain and does not activate Arp2/3 (and not nucleate actin in flies) ^35^. NPFs themselves also require activation that prevents intra- or inter-molecular shielding of their VCA domains. WASp and WAVE are autoinhibited and are activated upon binding of Cdc42 and PIP₂ (phosphatidylinositol 4,5-bisphosphate) or Rac1, respectively ^36–39^. WASH can be activated by RhoA, but is not autoinhibited. Rather, its VCA domain is spatially constrained by the multiprotein WASH complex, mutations in which result in neurological pathologies including intellectual disability and Alzheimer ^40–44^. It is comprised of FAM21, Strumpellin (aka. WASHC4), SWIP, and CCDC53 and prevents uncontrolled actin nucleation at endosomal membranes ^45–47^.

Intriguingly, our data show that overexpression of the NPFs *WASp*, *Wash* and *SCAR* alters the kinetics of e-MI from 25 h to 4 h of exposure to stress. This expedited e-MI is dependent on a stress trigger and the ESCRT machinery and can also be elicited by overexpression of the Arp3 subunit of the Arp2/3 complex. Interestingly we observed that among the *Drosophila* NPFs, only Wash is required for physiological induction of e-MI. Mechanistically, Wash activates the Arp2/3 complex to induce e-MI, indicating a specific requirement for actin branching at the late endosomal surface. Wash activates e-MI as part of the WASH complex, but independent of retromer that can recruit the WASH complex to early endosomes. Together, our findings demonstrate that forced expression of NPFs is sufficient to reprogram the temporal dynamics of e-MI and identify a critical role for NPF-mediated regulation of actin at late endosomes in controlling e-MI. We thus have identified a novel physiological mechanism regulating e-MI that is well positioned to be used to manipulate e-MI in vivo.

## Results

### The actin nucleation promoting factor WASp changes the temporal induction of e-MI

Constitutive e-MI controls the stability of proteins such as WASp (Wiskott Aldrich syndrome protein) and Comatose/NSF1 (N-ethylmaleimide-Sensitive Factor 1) that promotes synaptic vesicle turnover at 3^rd^ instar neuromuscular junctions ^23^. We wondered whether Comt or WASp are e-MI substrates in the FB which is a tissue commonly used to assess autophagy ^22, 25, 26^. If so, it is particularly interesting to determine whether they also behave as constitutive e-MI substrates or rather are substrates of stress induced e-MI. We thus generated transgenic flies expressing photoactivatable mCherry (PA-mC)-tagged fusions of WASp and Comt analogous to our standard e-MI sensor based on an artificial fusion of the KFERQ motif of RNAseA to PA-mC ^22^. These sensors then were expressed under the control of a FB specific Gal4 driver line (*cg-Gal4*) ^48^ and photoactivated in early 3^rd^ instar larvae that were then transferred either to 20% sucrose (for starvation) or to 20% sucrose solution supplemented with heat-killed yeast (fed) for 25h ^22^. Both Comt and WASp sensors formed puncta upon prolonged starvation, while remaining diffuse in the cytoplasm under fed conditions (Fig. 1A–D, quantified in 1E; see Table S1 for exact genotypes). To verify that the sensor puncta reflect e-MI induction, we assessed if their formation depended on the ESCRT machinery. Indeed removing the ESCRT II subunit *Vps25* or the ESCRT III component *Vps32/shrub* using the FRT-FLP technique in genetically mosaic animals ^49^ prevented the formation of mCherry puncta in cell-autonomously in mutant cells expressing PA-mC-Comt or PA-mC-WASp (Fig. 1I-K; compare homozygous mutant cells lacking GFP expression with GFP expressing control cells). The starvation and ESCRT dependence of puncta formation indicates that, in the FB, WASp and Comt are e-MI substrates, but only under conditions of starvation induced stress.

**Figure 1.**
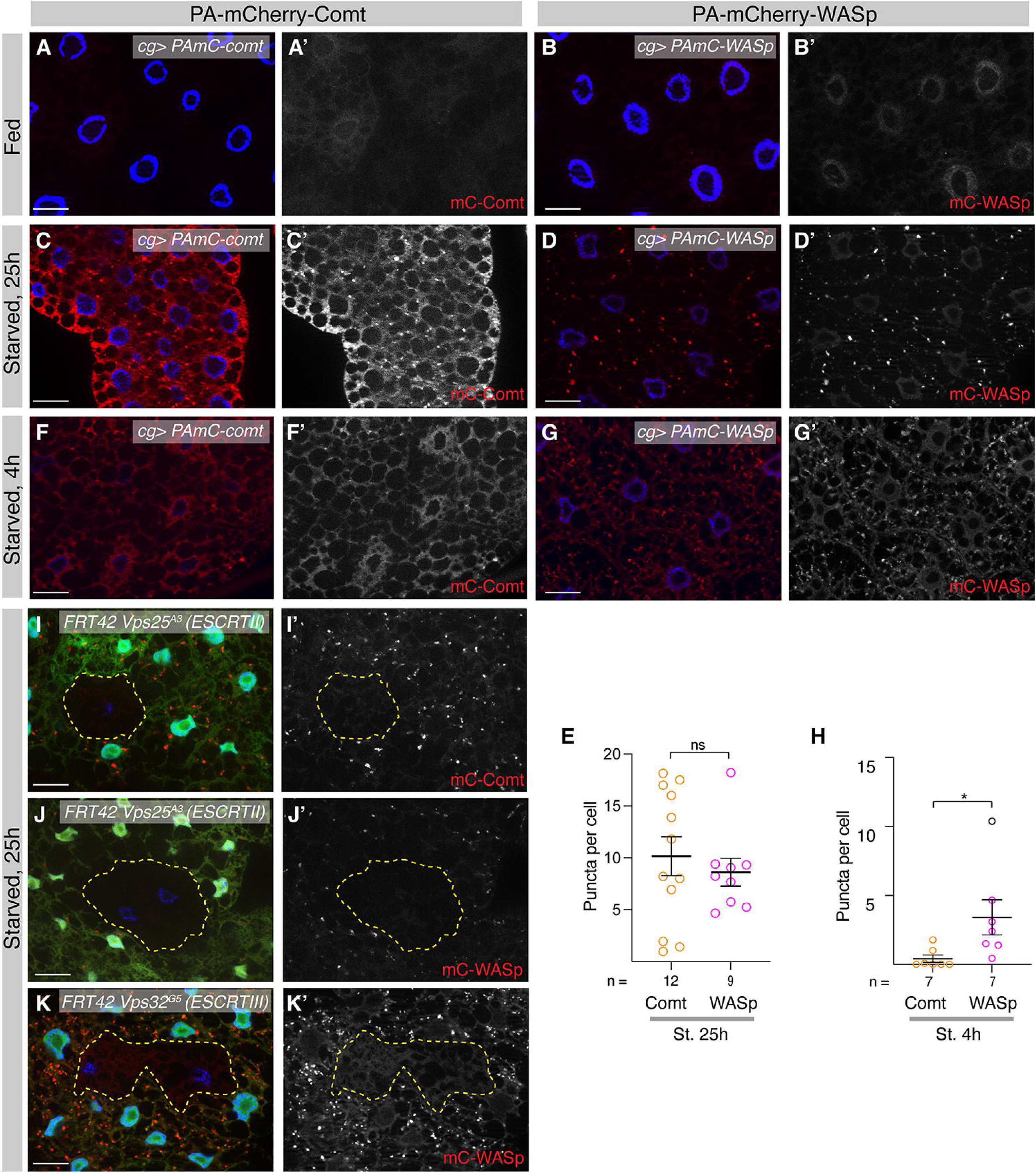
PA-mC-WASp alters the timing of e-MI. **(A-D)** Under fed conditions PAmC-Comt (A) and PAmC-WASp (B) e-MI substrates are diffusely distributed in 3^rd^ instar larval fat body cells 25h after photoactivation. Upon 25 starvation, PAmC-Comt (C) and PAmC-WASp (D) show punctate patterns typical for e-MI substrates. **(E)** Quantification of puncta number per cell at 25hps for PAmC-Comt and PAmC-WASp. **(F, G)** Compared to PAmC-Comt (F), PAmC-WASp (G) forms puncta already at 4hps. **(H)** Quantification of sensor puncta at 4hps. **(I-K)** PAmC-Comt puncta formation is dependent on the ESCRT II component Vps25 (I) and PAmC-WASp puncta formation requires the ESCRT II component Vps25 (J) and the ESCRT III component Vps32/Shrb (K). Homozygous mutant cells are marked by the absence of GFP and are outlined by a dotted yellow line; WT and heterozygous cells express (nuclear) GFP. Monochromatic images show the indicated channels. Nuclei are in blue. Scale bars: 20 µm. Student t-tests; * p< 0.05; ns: not significant; n: fields of view.

A striking feature of starvation-induced eMI is its delayed activation (12-25h) compared to macroautophagy (1-4h) ^22, 24^. To our surprise, we found that upon expression of the WASp, but not the Comt sensor, e-MI was already induced after short-term (4 h) starvation (Figure 1F, G; quantified in Fig. 1H), suggesting that elevated levels of PA-mC-WASp in FB cells result in earlier e-MI or that WASp could localize to e.g. autophagosomes. To test this, we knocked down *Atg7* by RNAi with a line that had previously been shown to inhibit MA in the *Drosophila* FB ^22^. Blocking MA did not prevent PA-mC-WASp from forming puncta upon 4h of starvation (quantified in Fig. S1A). To confirm that the WASp puncta reflect expedited e-MI, we generated homozygous mutant clones in the fat body for *Vps32^G^*^5^ (ESCRT-III) in which PA-mC-WASp sensor puncta failed to from 4h post starvation (Fig. 2G), suggesting that WASp expression indeed changes the time course of e-MI induction. Consistently, WASp induced e-MI also requires the function of Atg1 known to mediate starvation induction of e-MI and MA upon TOR inhibition (quantified in Fig. S1A) ^22, 25, 26^.

**Figure 2.**
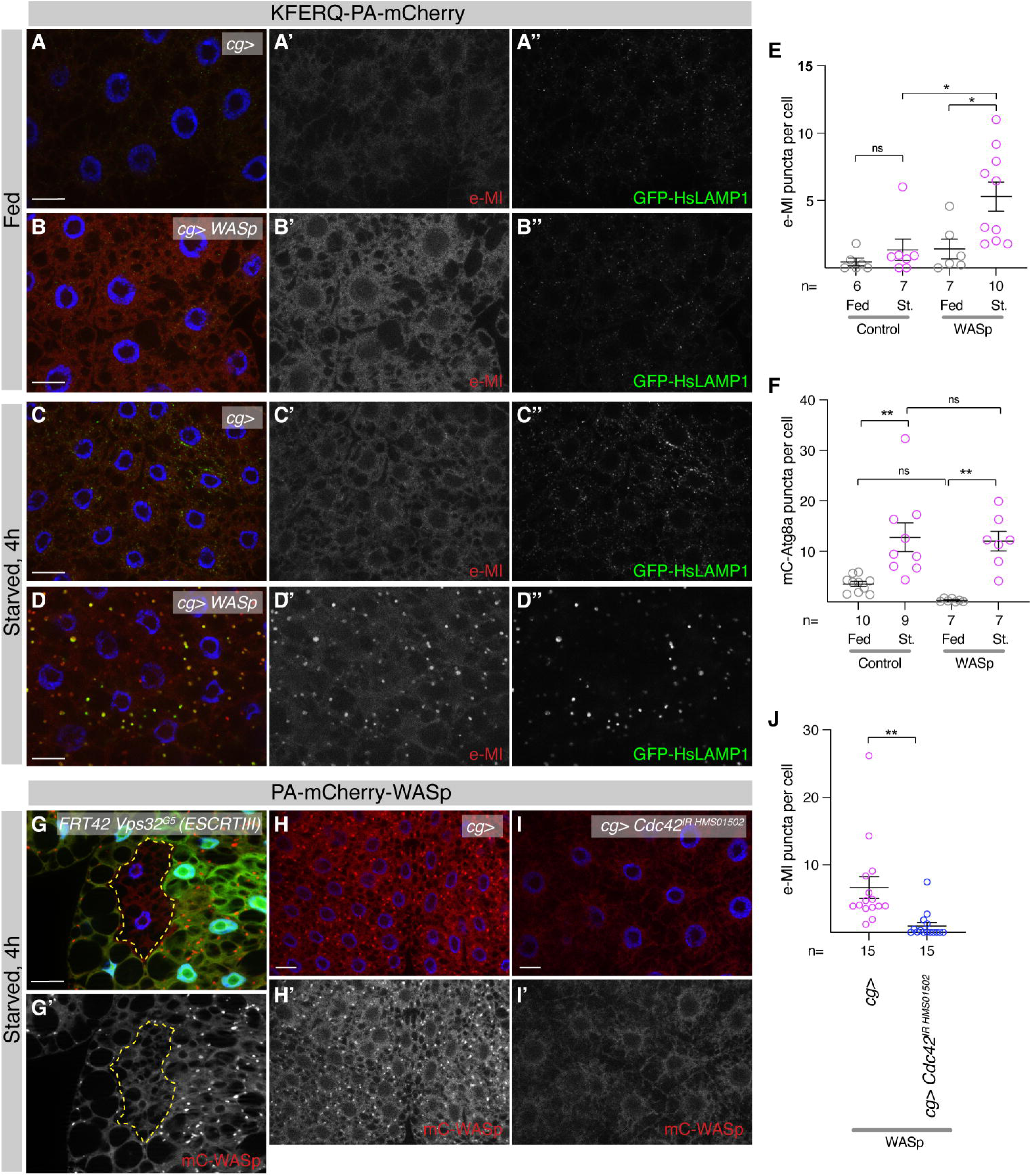
WASp induced expedited e-MI is Cdc42 dependent. **(A-E)** Under Fed conditions, both in control (A; *cg-Gal4*) and upon expression of WASp (B), the regular KFERQ-PAmC e-MI sensor is diffusely localized in 3^rd^ instar FB cells at 4h. However, while the e-MI sensor in the control (C) remains diffuse at 4hps, WASp overexpressing cells (D) show a strong e-MI response. e-MI puncta overlap with the (late)-endolysomal marker HsGFP-LAMP1 (D’’). (E) Quantification e-MI response under indicated conditions. **(F)** Quantification of mCherry–Atg8a puncta reflecting autophagosomes and autolysosomes at 4hps, the peak time of MA induction under control conditions and upon WASp expression. WASp expression has no effect on Atg8a puncta. **(G)** WASp induced early e-MI is ESCRT dependent, as *Vps32^G^* mutant cells (outlined by dotted yellow line and lacking the GFP marker) fail to from PAmC-WASp sensor puncta. **(H-J)** Compared to WASp expression alone (H), RNAi mediated knockdown of *Cdc42* prevents early e-MI induction by WASp (I), showing that it requires physiological activation of WASp. (J) Quantification of e-MI puncta per cell in control and *Cdc42RNAi*. Monochromatic images show the indicated channels. Nuclei are in blue. Scale bars: 20 µm. One-way ANOVAS (Tukey) p<0.01 (E) and p<0.0001 (F); J: Student t-test; * p< 0.05; ** p< 0.01; ns: not significant; n: fields of view.

To rule out the possibility that expedited e-MI is an artifact due to the expression of WASp as mCherry fusion protein, we expressed wildtype, untagged WASp in the FB in the presence of the regular KFERQ-PA-mC e-MI sensor and a (endo)lysosomal Marker (GFP-HsLAMP1). We found that WASp is able to induce e-MI puncta upon 4h of starvation (Fig. 2A-D; quantified in 2E). As expected, the puncta overlap with late endosomes and lysosomes (Fig. 2D) ^22^. Importantly, compared to control, WASp overexpression did not alter the number of autophagosomes and autolysosomes marked by mCherry-Atg8a ^27^, the *Drosophila* homolog of LC3 (quantified in Fig. 2F). Physiologically, WASp is maintained in an autoinhibited conformation until it is activated by the small GTPase Cdc42 ^50, 51^. To test whether WASp induced expedited e-MI requires Cdc42, *Cdc42* was depleted by RNAi-knockdown specifically in the FB, which significantly reduced mC-WASp induced early e-MI (Fig. 2H,I; quantified in 2J). Therefore, in addition to a starvation signal, canonical upstream activation of WASp by Cdc42 is required for it to accelerate e-MI.

We next tested whether WASp is required for stress induced e-MI. To assess the cell-autonomous requirement of WASp in e-MI, we generated mosaics of strong hypomorphic *WASp*^1^ allele ^52^ in the FB. We observed that the number of sensor puncta induced in *WASp*^1^ mutant cells was comparable with surrounding wild type cells (compare the *WASp*^1^ mutant cells lacking GFP with their neighbors in Fig. 3A; quantification in 3B). Consistent with this result we found that knockdown of *Cdc42* showed no effect on e-MI after 25h of starvation (Fig S2A-D, quantified in S2E). These results show that although WASp is sufficient to expedite e-MI in a manner dependent on Cdc42, neither WASp nor Cdc42 is required for e-MI.

**Figure 3.**
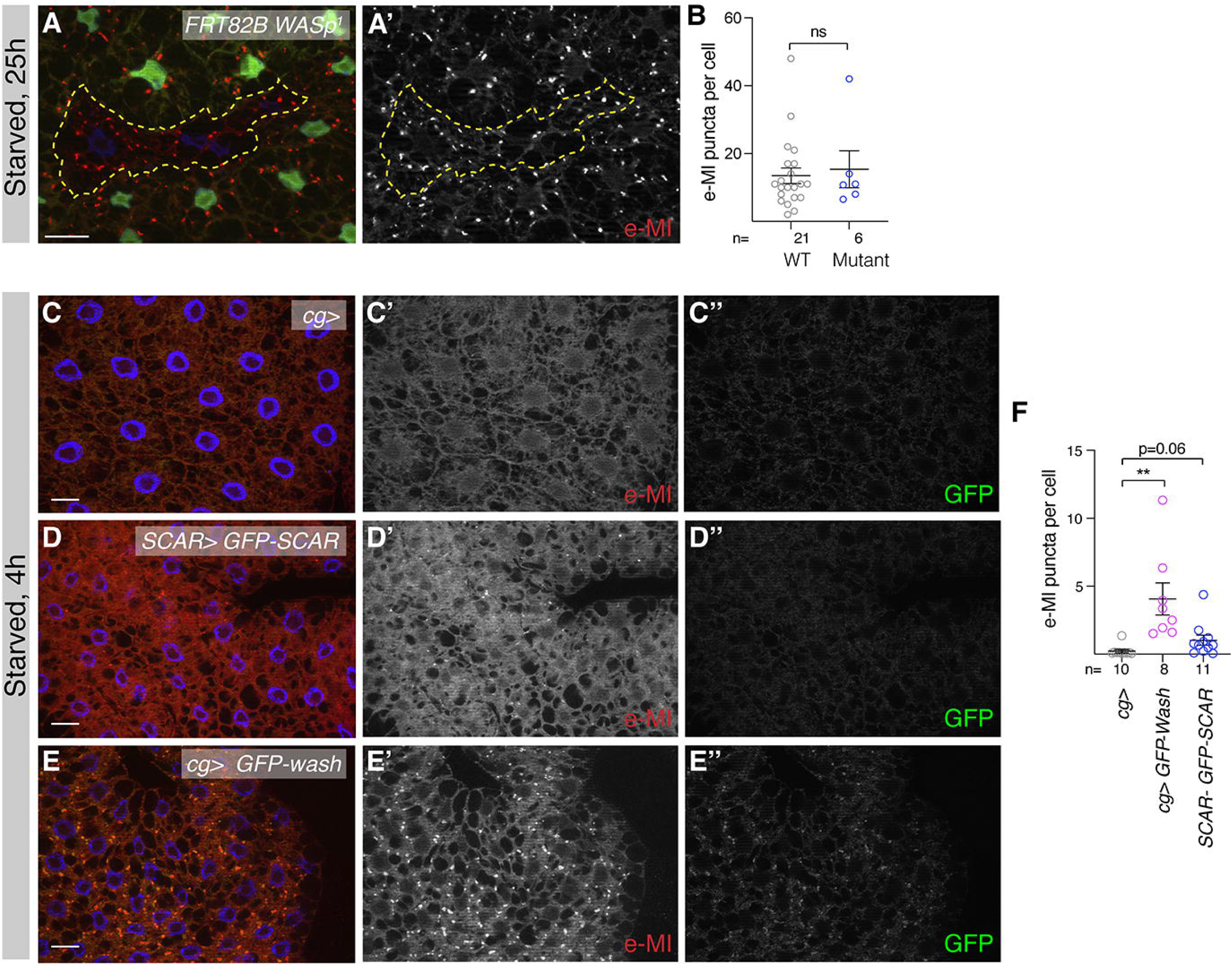
Wash and SCAR also accelerate e-MI induction upon starvation. **(A, B)** WASp is not required for e-MI, as *WASp*^1^ mutant cells lack e-MI sensor puncta at 25hps (A; compare mutant cells outlined with yellow dotted line and lacking GFP with surrounding WT cells). (B) Quantification of e-MI puncta in wild type and *WASp*^1^ mutant cells. **(C-F)** Compared to KFERQ-PAmC sensor only (C), expression of *GFP-SCAR* under control of its own promoter (D) shows a tendency towards e-MI activation. Expression of *GFP-Wash* (E) induces a robust activation of e-MI at 4hps. Note the overlap between GFP-Wash and the (late)endolysosomes marked by the e-MI sensor. (F) Quantification of e-MI response at 4hps in FB of indicated genotypes. Monochromatic images show the indicated channels. Nuclei are in blue. Scale bars: 20 µm. Student t tests; ** p< 0.01; ns: not significant; n: number of cells in (B) and fields of view in (F).

### Overexpression of Wash and SCAR also can expedite e-MI

Based on WASp being sufficient to alter the timing of, but not required for e-MI, we wondered if NPFs are redundant for e-MI induction, in particular as they are known to all nucleate actin *in vitro* and to have overlapping as well as distinct functions ^53–55^. The *Drosophila* genome codes for two additional Arp2/3-activating NPFs, Wash and SCAR. We thus assessed whether Wash and SCAR expression can also induce e-MI upon starvation for 4h. A transgene expressing *GFP*-*SCAR* driven by its endogenous promoter shows a tendency towards accelerated e-MI activation (Fig. 3D; quantified in 3F), although the increase associated with GFP-SCAR did not reach statistical significance (p=0.06), likely due to the comparatively lower expression level of the genomic *SCAR* construct. In contrast, FB-specific expression of Wash robustly induces e-MI sensor puncta (Fig. 3E; quantified in 3F).

To determine whether the NPF response is specific for starvation, we exposed 3^rd^ instar larvae co-expressing Wash and the e-MI sensor in the fat body to ROS and genotoxic stress, which previous work from our lab showed to induce e-MI ^24^. Feeding larvae etoposide (Eto), inducing DNA double strand breaks, or paraquat to induce ROS for 4h induced a robust early activation of e-MI provided that Wash is overexpressed (quantified in Fig. S3). This result indicates that NPF-induced changed timing of e-MI follows still requires a physiological trigger, but is not restricted to activation by starvation.

### Wash is required for e-MI

Since NPFs are sufficient to drive early e-MI and WASp is not required for e-MI, we next assessed whether Wash and SCAR are required or redundant with WASp. RNAi-mediated knockdown of *Wash* and *Scar* in the fat body shows that depletion of *Wash*, but not *Scar*, resulted in a significant reduction in e-MI sensor puncta (Fig. 4A–C’’; quantified in Fig. 4D; note that SCAR^IR^ ^HMC03361^ was successfully used to inactivate *SCAR* during oogenesis ^56^). Importantly, two independent, non-overlapping RNAi lines targeting *Wash* consistently showed decreased e-MI puncta upon 25h of starvation, supporting a specific requirement for Wash in e-MI activation (quantified in Fig. 4E). The physiological functions of NPFs depend on their distinct subcellular localization. WASp and SCAR predominantly function at the plasma membrane, where they regulate Arp2/3-dependent actin assembly, while Wash is known to localize to early and late endosomes ^57^. Consistently, we see a strong co-localization of Wash-GFP and KFERQ-PA-mC reporter puncta suggesting that Wash localizes to late endosomes and lysosomes (Fig. 3E-E’’)^22^.

**Figure 4.**
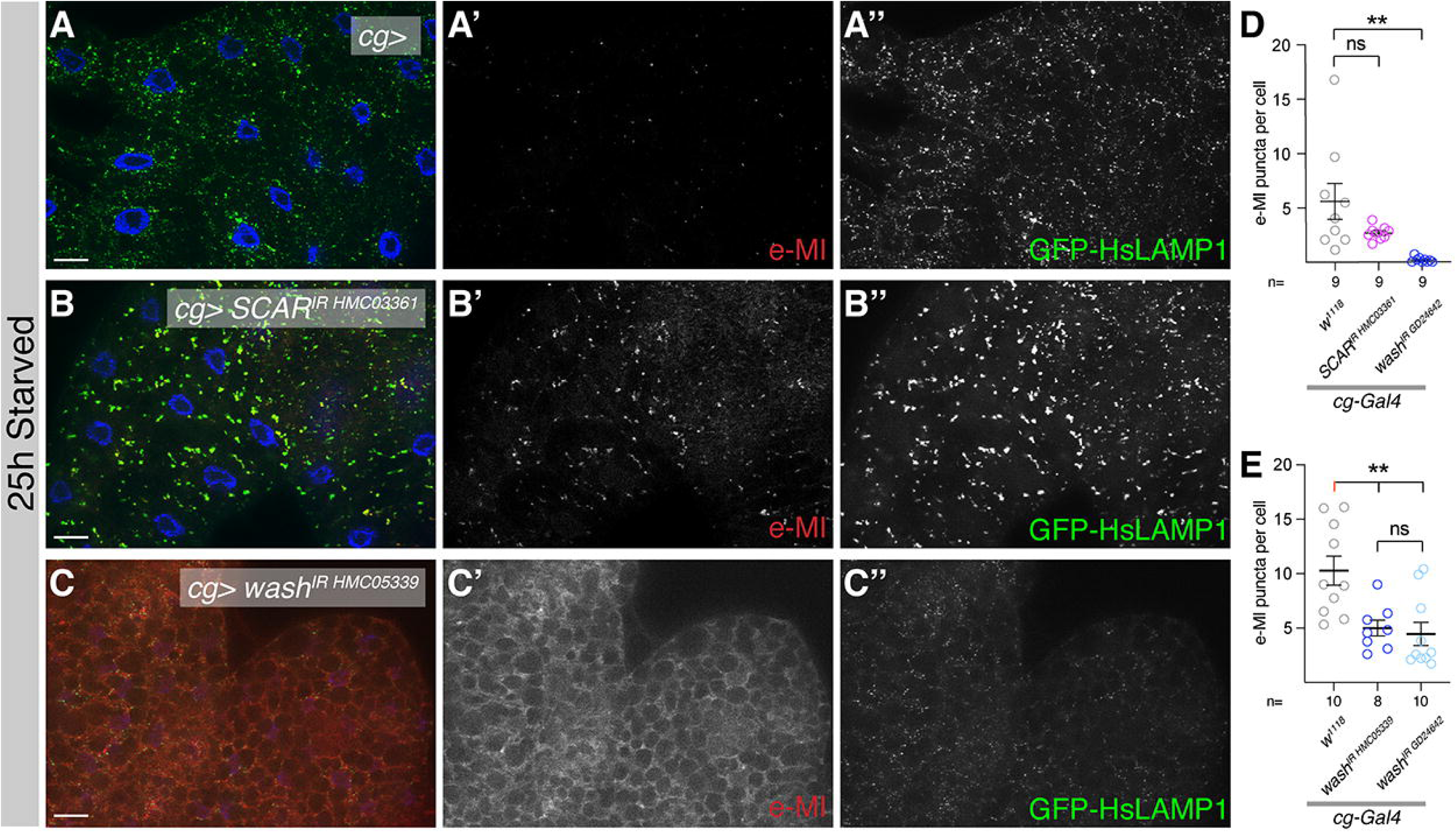
Wash is required for e-MI. Upon starvation for 25h, e-MI is induced in control FB **(A)** and in FB in which *SCAR* is knocked down by RNAi **(B)**. In contrast, knockdown of *wash* prevents e-MI activation upon prolonged starvation **(C)**, showing that *wash* is essential for e-MI and that NPFs are not redundant for this function. **(D, E)** Quantification of e-MI responses in indicated genotypes at 25hps. Note that the two *wash* RNAi lines have non-overlapping targets, thus excluding off-target effects. Monochromatic images show the indicated channels. Scale bars: 20 µm. One-way ANOVAS (Dunnett) p<0.01; ** p< 0.01; ns: not significant; n: fields of view.

NPF activation often involves the Rho family GTPases and Rho1, Rac, and Cdc42 that have distinct specificities for the regulation of WASH, SCAR, and WASp. Wash binds to active, GTP-bound Rho1, which activates Wash by overcoming inhibition by Spire during *Drosophila* oogenesis ^58^. We thus clonally expressed dominant-negative Rho1^N19^ using the flip-on system in which expressing cells are marked with GFP ^59^. Dominant-negative Rho1 did not affect number of e-MI reporter puncta compared to surrounding wild-type tissue (Fig. S4A). The same result was obtained in *Rho1^M^*^1^ mutant cells in mosaic analysis (S4B). We also generated flip-on clones expressing dominant negative *Rac1^N^*^17^ (marked with GFP) that did not change e-MI sensor puncta (Fig. S4C) at 25hps. Collectively, these data indicate the Rho-family GTPases Rho1, Rac, and Cdc42 are not required for e-MI, suggesting that Wash is activated in a distinct manner.

### Wash acts as part of the WASH complex, but independent of retromer to activate Arp2/3 and promote e-MI

We then addressed how Wash regulates e-MI and first asked whether the effect of NPFs on e-MI functions through activation of the Arp2/3 complex and thus branched actin. To test this possibility, we co-expressed the GFP tagged Arp3, a subunit of Arp2/3 complex, in fat body of *Drosophila* along with KFERQ-PA-mC reporter to monitor e-MI. Expression of GFP-Arp3 in the FB results in induction of KFERQ-PA-mC reporter puncta at 4 hps (Fig. 5A; quantified in 5E), suggesting that activation of Arp2/3 is sufficient to expedite e-MI upon stress exposure.

**Figure 5.**
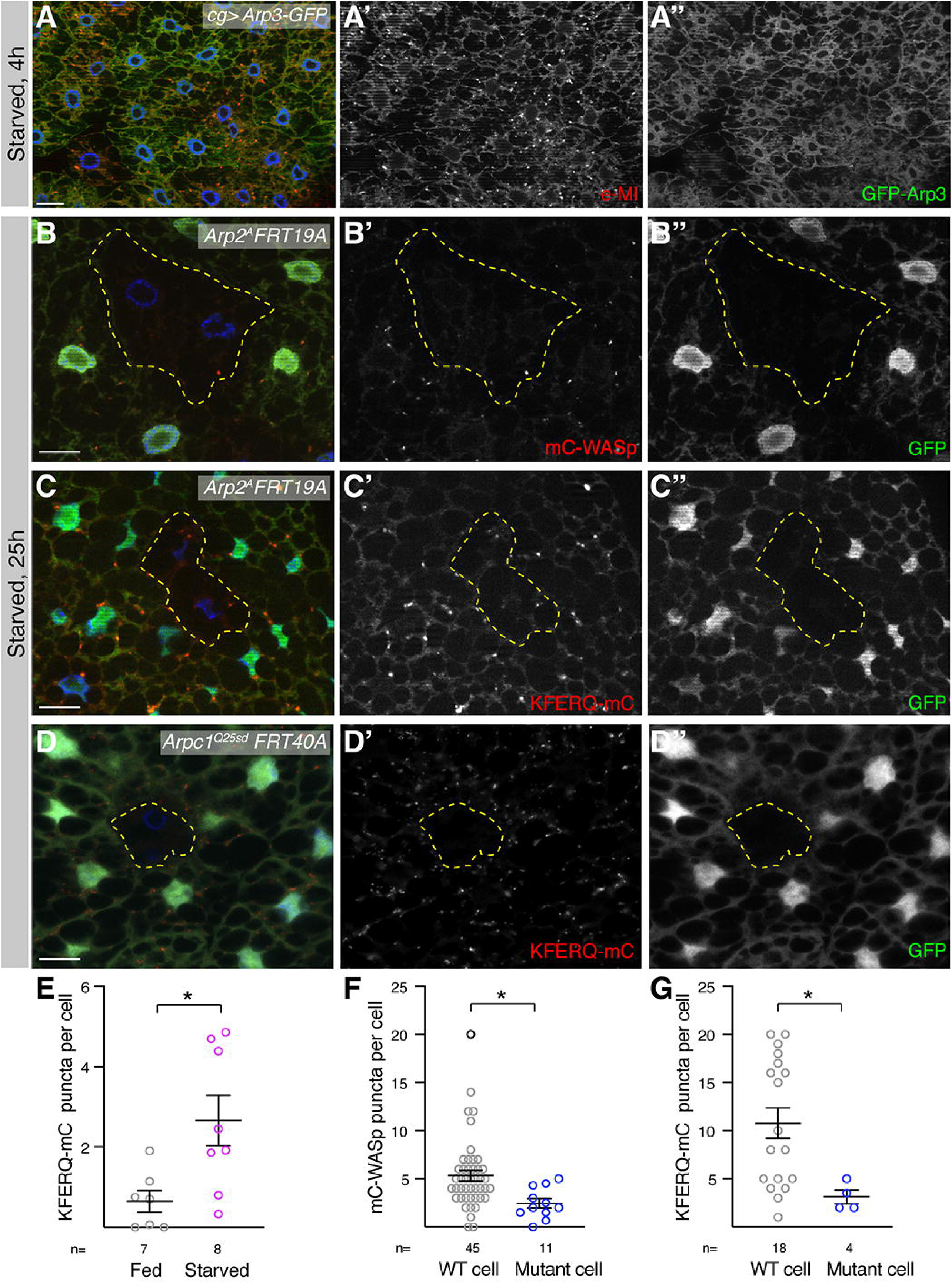
NPFs act through Arp2/3 to activate e-MI. **(A)** Upon 4h of starvation, fat body expression of *Arp3-GFP* is sufficient to induce e-MI as assessed by KFERQ-PAmC sensor puncta formation (quantified in E). **(B-C)** Mosaic analyses shows that cells mutant for *Arp2^A^* (lacking GFP expression and surrounded by a dotted yellow line) fail to show an e-MI response assessed using the PAmC-WASp (B; quantified in F) or regular e-MI sensor (C; quantified in G), showing that the NPFs act though the Arp2/3 complex to regulate e-MI. **(D)** The *Arpc1^Q25SD^* null mutant cells also fail to form KFERQ-PAmC sensor puncta compared to surrounding wild type cells. Note that only a very small number of mutant cells was obtained. **(E)** Quantification of early e-MI induced by expression of Arp3-GFP. **(F, G)** Quantification of e-MI in WT and *Arp2* mutant cells measured by PAmC-WASp (F) and KFERQ-PAmC (G), respectively. Monochromatic images show the indicated channels. Nuclei are in blue. Scale bars: 20 µm. Student t tests; * p< 0.05; n: number of fields of view in (A) and cells in (B, C).

To test whether NPF-driven recruitment of the Arp2/3 complex also is required for stress induced e-MI, we generated mutant clones for *Arp2^A^*, a likely hypomorphic allele and the null allele *ArpC1^Q^*^25s*d*^ ^60^, two subunits of the Arp2/3 complex in otherwise heterozygous background. Both KFERQ-PA-mC reporter and PA-mC-WASp sensor puncta formation was significantly decreased in *Arp2^A^* mutant cells when compared to wild type cells at 25 hps (Fig. 5B, C; quantified in Fig. 5F and 5G, respectively). Similarly, we see a lack of KFERQ-PA-mC puncta *ArpC1* mutant clones (Fig. 5D; note that due to the limited number of recovered clones, this analysis was based on a sample size of only n=3). These results show that Arp2/3 recruitment is critical for e-MI activation. Multiple NPF family members have been shown to redirect Arp2/3 toward distinct stress-induced functions including autophagy ^5, 61–64^. We therefore tested if Arp2/3 machinery in *Drosophila* is also required for MA and made *Arp2* mutant clones in the presence of a *mCherry-Atg8a* marking autophagosomes and autolysosomes. *Arp2* mutant cells showed no change in mCherry-Atg8a puncta induced upon 4 h of starvation, the peak time of MA (Fig. S1C, quantified in S1D) suggesting that Arp2/3 requirement is specific to e-MI and Arp2/3 machinery is not required for MA initiation in the *Drosophila* FB.

In both mammals and *Drosophila*, Wash can exist as part of a multi-subunit WASH regulatory complex that includes FAM21, Strumpellin, SWIP, and CCDC53, and keeps Wash inactive and functions to regulate and spatially restrict actin nucleation at endosomal membranes ^45–47^. The WASH complex can be recruited to endosomes via FAM21 binding to the Vps35 subunit of the retromer complex. Retromer, recruited to membranes by its Syntaxin 3 (Snx3) subunit, controls recycling of cargo proteins in conjunction with the WASH complex from (early) endosomes back to either the Golgi apparatus or the plasma membrane ^65^. To test whether other WASH complex components participate in e-MI, we knocked down *FAM21* with RNAi in the FB and observed significant depletion of e-MI reporter puncta compared to Gal4-driver only control at 25 hps (Compare Fig. 6A and 6B; quantified in 6C; RNAi efficiency quantified in Fig. S5), suggesting that the WASH complex is required for e-MI. This led us to examine the involvement of retromer components in e-MI. While we find a partial reduction of e-MI upon knockdown of Snx3 (Fig. 6C; Fig. S5 for RNAi efficiency), loss of core retromer subunit *Vps35* in null mutant cells ^66^ did not alter KFERQ-PA-mC reporter puncta formation upon 25h of starvation (Fig. 6D; quantified in Fig. 6E), showing that retromer is not required for e-MI. To address whether expedited e-MI activation induced by Wash overexpression altered the endolysosomal system, we stained FB overexpressing Wash and the e-MI sensor for the late endosomal and lysosomal markers Rab7 and HsLAMP1-GFP. Upon starvation, Wash overexpression caused a strong expansion of Rab7-positive structures, which co-localized with HsLAMP1-GFP (Fig. S6A, B; quantified in S6C).

**Figure 6.**
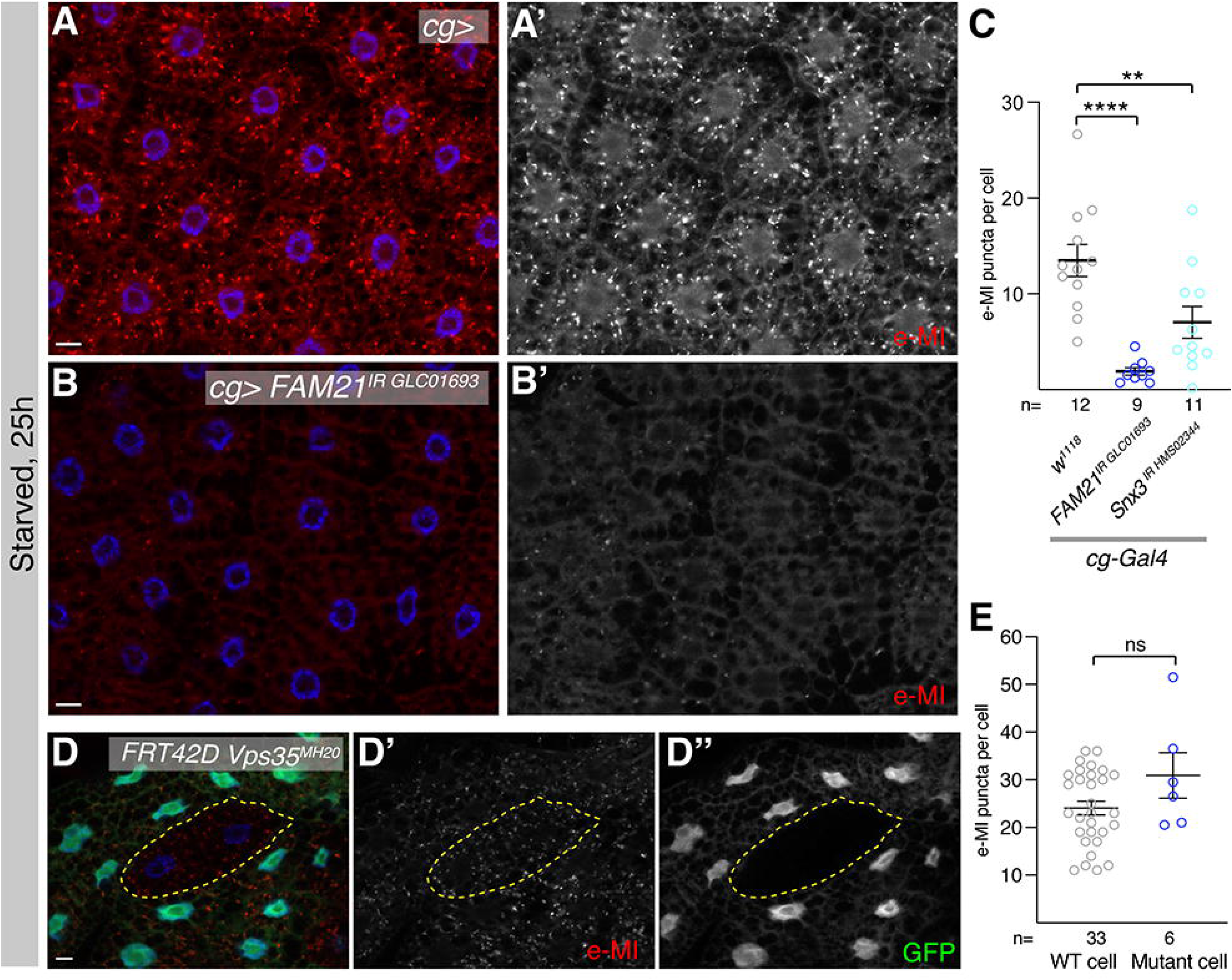
Wash acts as part of the WASH complex, but independent of retromer. **(A, B)** Compared to sensor only control (A), RNAi mediated knockdown of the WASH complex component *FAM21* in the fat body of 3^rd^ instar larvae prevents e-MI at 25hps (B). **(C)** Quantification of puncta number per cell in *FAM21* and *Snx3* RNAi at 25hps. **(D)** Cells mutant for *Vps35^MH^*^20^, a core subunit of retromer, showed no difference in e-MI activation after prolonged starvation, indicating that retromer is not required for e-MI. Mutant clones are marked by the absence of GFP and outlined by a dotted yellow line. **(E)** Quantification of e-MI activity at 25hps, in WT and *Vps35^MH^*^20^ mutant cells. Monochromatic images show the indicated channels. Nuclei are in blue. Scale bars: 20 µm. (C): One-way ANOVA (Tukey) p < 0.0001; (E): Student t-test; ** p< 0.01; *** p< 0.001; n: number of fields of view in (C) and cells in (E).

Overall, our results suggest that the WASH complex, mutation of which have been linked to neurodegenerative diseases ^40–44^, is required for endosomal microautophagy. Intriguingly, this function of WASH is independent of the core retromer component Vps35 and may thus represent a novel mechanism of how WASH regulates late endosomal functions that are associated with degradative trafficking rather than endosomal recycling. Despite this distinct mode of action, the WASH function ultimately seems to involve the formation of branched actin networks, as Arp2/3 activity is necessary and sufficient for e-MI modulation.

## Discussion

Here, we report a novel, essential function of Wash, an actin nucleation-promoting factor, which activates the Arp2/3 complex to generate branched actin filament networks, in the regulation e-MI. We showed that forced expression of a NPF accelerates e-MI activation upon shorter stress exposure. While several NPF family members can induce e-MI, they do not act redundantly, as Wash specifically is required for e-MI under physiological conditions in the *Drosophila* larval FB. For e-MI, Wash acts via the Arp2/3 complex, suggesting that e-MI activation requires the formation of branched actin at the late endosome. Overall, we report that NPFs mediated actin cytoskeleton branching mediates stressed induced activation of e-MI. Intriguingly, while Wash acts as part of the WASH complex, its function is independent of a key component of the retromer complex, which is known to recruit the WASH complex to early endosomes for its function in protein recycling ^65^. Despite NPFs or Arp3 being sufficient to expedite e-MI, they do not uncouple it from a stress trigger.

In mammals, the WASH complex is ubiquitously expressed and mutations in WASH complex components result in neurological pathologies including intellectual disability and Alzheimer ^40–44^. In its best-known function, Wash acts at early endosomes, where the 5-subunit WASH complex is recruited by retromer to promote actin remodeling dependent cargo recycling back to the Golgi or the plasma membrane, thereby preventing cargo from lysosomal degradation ^45, 65, 67^. While depletion of the WASH complex member FAM21, which interacts with retromer, blocked e-MI activation in the fat body, the classical recycling pathway is not required for e-MI, as loss of *Vps35*, a core component of retromer, did not alter stress-induced e-MI sensor puncta formation. These findings indicate that the role of Wash in e-MI activation reflects a novel, non-canonical function associated with degradative trafficking rather than endosomal recycling. This interpretation is consistent with previous reports showing that retromer-dependent recycling becomes less prominent during starvation, when cellular trafficking is redirected towards degradation ^68^. It also is worth noting that silencing *WASH* in fibroblasts reduces trafficking of endocytosed EGFR to LAMP1 positive endolysosomes ^69^ and *VPS35* knock-out in HeLa cells reduces basal EGFR levels, which would be consistent with EGFR taking a degradative route in the absence of retromer ^70^ and WASH being involved in the decision over recycling versus degradation.

Our data are consistent with a model in which the Wash complex works at late endosomes to promote e-MI. Consistently, GFP-Wash and e-MI sensor puncta, the latter known to strongly overlap with the late endosomal marker Rab7 and the endolysosomal marker HsLAMP1-GFP ^22^, colocalize upon starvation (Fig. 3E). Furthermore, Wash expression results in expansion of Rab7 positive puncta upon short starvation while having no effect under basal conditions (Fig. S6). Under stress, Wash could thus result in actin remodeling at late endosomes which may reorganize the Rab7 compartment to activate or stimulate e-MI.

How does Wash alter actin to affect e-MI? Wash not only can induce Arp2/3 mediated actin branching, but also can bundle actin fibers (and microtubuli) *in vitro*, a function that is regulated by RhoA *in vitro* and *in vivo* ^58^. As we find no (unique) involvement of Rho GTPases in e-MI, the role of Wash in e-MI likely does not involve actin bundling but rather branched actin formation, consistent with Arp2/3’s most prominent function on early endosomes and also supported by the fact that Arp2/3 antagonizes actin bundling by Wash *in vitro* ^58^. Wash can also be activated by phosphorylation of Bruton Tyrosine kinase (BTK) to favor the formation of endosomal actin at the expense of cortical actin, which in *Drosophila* regulates the timing of tracheal clearing necessary for breathing ^71^. However, RNAi mediated knockdown of *Btk* did not prevent e-MI nor was Wash^Y261^, mimicking Btk activation ^71^, able to promote e-MI at 25hps (not shown), suggesting that Btk is not activating Wash for e-MI. Future experiments are needed to identify the activator of Wash in e-MI.

In contrast to NPF activation, we found that the downstream execution of the e-MI process likely proceeds through a conserved molecular machinery as Wash, WASp, and likely SCAR overexpression all lead to activation of expedited e-MI (Figs. 2, 3). This also is directly supported by our finding where abolishing Arp2/3 function by mutation of *Arp2* and *ArpC1* impaired e-MI, while overexpression of Arp3 was sufficient to induce early e-MI upon starvation. The stress-dependence of e-MI is consistent with the emerging view of Arp2/3 as a conditional stress responder rather than a constitutively active effector. Under basal conditions, Arp2/3 activity is tightly constrained and spatially restricted to endocytic patches and kept in check by the limited availability of distinct NPFs. Multiple WASP-family nucleation-promoting factors have been shown to redirect Arp2/3 toward distinct stress-induced functions including autophagy, DNA repair, and membrane remodeling ^61–64^. Notably, while several NPFs are capable of engaging Arp2/3 in a stress context, our observations suggest that, physiologically, the specific effect on e-MI is driven by Wash coupling different upstream stress signals to a precise actin assembly to promote e-MI. The requirement for a stress trigger even in the context of NPF overexpression indicates that Arp2/3 activation alone is insufficient to drive this phenotype. Rather, stress appears to create a permissive cellular context potentially through remodeling of membrane composition, organelle dynamics, or cofactor availability ^72, 73^.

A role of NPFs and the Arp2/3 complex as a regulators of macroautophagy is also emerging, although existing research is not easily reconciled and NPF/Arp roles may depend on the context. WASp can promote autophagosome formation and autophagosome–lysosome fusion ^62, 63^}. WASH modulates ATG9 vesicle trafficking and regulates the Beclin1–PI3KC3 complex to control autophagy induction, and the WAVE/SCAR complex supports autophagosome biogenesis, consistent with a role for Arp2/3-dependent actin nucleation in this process ^61, 74^. In yeast, Arp2/3 also regulates Atg9p trafficking and ts-mutations in *Arp2* slow down MA and prevent autophagic degradation of peroxisomes (pexophagy) ^75^. In contrast, inducible knockout of Arp2 in mouse ear fibroblasts doesn’t block autophagosome formation, but rather leads to an increase in autolysosomes due to defects in autolysosome reformation ^76^. It also has been reported recently that MA can be inhibited by WNT2B mediated WASHC5/SWIP function ^77^. In the fly FB, we find no change in mC-Atg8 puncta in *Arp2* mutant cells (Fig. S1 C,D), suggesting that neither autophagosome formation nor autolysosome reformation (ALR) is defective, as the former would lead to a reduction and the latter to an increase in mC-Atg8 puncta (this reporter cannot distinguish between autophagosomes or autolysosomes).

Loss of NPF function compromises the endo-lysosomal integrity that neurons depend on for protein and organelle clearance ^78^ and disruption of the WASH complex produces endo-lysosomal defects with cognitive and motor consequences in both mice and humans ^79^. On the other hand, the endo-lysosomal compartment serves as a convergence point for the various forms of autophagy and is broadly compromised across neurodegenerative diseases including Alzheimer’s, Parkinson’s, and frontotemporal dementia, and thus represents a shared pathological node rather than a disease-specific defects ^10^. Our study identifying Arp2/3 activity at the endo-lysosome as a previously unrecognized regulator of e-MI, thus opens the possibility of targeting branched actin assembly to enhance e-MI in neurodegenerative disease and promote aggregate clearance in the future.

## Methods

### Drosophila strains

Flies were maintained on standard cornmeal medium at 25°C unless otherwise indicated. The following fly lines were used in this study (exact genotypes are listed in Table S1): *cg-Gal4, hsflp; cg-Gal4 FRT42D UAS-GFPnls, hsflp; r4-Gal4 FRT82B UAS -GFPnls* were kind gifts of Dr. T. Neufeld. *hsFlp act>CD2>Gal4 UAS-GFP, yw hsflp; UAS-mCherry-Atg8a* ^27, 80^*, atg8a>3xmCherry Atg8a* ^81^ were kind gifts of Dr. G. Juhasz. *UAS-GFP-HsLAMP1/CyO* was a gift from T. E. Rusten ^25, 82^, *UAS-Rho1^N^*^19–4.3^*, FRT Rho1^M^*^1^ were gifts of Dr. M Mlodzik (Mount Sinai School of Medicine, NY). UAS-wash was a kind gift of V. Tsarouhas (Stockholm University, Sweden) ^71^.

Additional lines used were: *r4-KFERQ-PAmCherry#8/ TM6B* ^24^; *FRT42D Vps25^A^*^3^*/CyO twist-Gal4-UAS-GFP, FRT42D Vps32^G^*^5^ *(shrub) /CyO twist-Gal4-UAS-GFP* ^83, 84^*, UASp-GFP-wash (*BL-81639), *SCAR-GFP-SCAR (*BL-94927) ^58^*, UAS-WASp* (BL-39724) ^52^, *FRT42D Vps35^MH^*^20 66^, *UASp-Arp3.GFP* (BL-39721), *UAS-Rac1^N^*^1–7–17^ (BL-6292) ^85^, *Arp2^A^ FRT19A (BL-52315)* ^86^, *Arpc1^Q^*^25s*d*^ *FRT40A (BL-9137)* ^60^. *WASp*^1^*/TM6B* (BL-51657) ^52^ was recombined onto FRT82B as described ^49^ and selected on food supplemented with 0.5 mg/mL G418 (Gibco). Recombinants were confirmed by PCR using primers WASp1-rec (Table S2). The following RNAi lines were used: *VDRC-GD16133* (*Atg1*), *VDRC-GD45558* (*Atg7*), *VDRC-GD24642* and *HMC05339* (BL# 62866; both *wash*), *HMS01502* (BL-35756; *Cdc42*); GLC01693 (BL-50571; *FAM21*), *HMS02344* (Bl-50571; *Snx3*), *HMC03361*(BL-51803 ;*SCAR*).

### Molecular biology and transgenic flies

The Gateway cloning compatible, photoactivable mCherry reporter containing vector (pTpaCW) was generated by modifying pTFW (a UAST-based Gateway vector ^87^). PAmCherry was amplified using the PAmCherry primers listed in Table S2. The PCR product was digested with AgeI and AvrII, and ligated into the corresponding sites of the pTFW to generate the pTpaCW vector. The open reading frame of *WASp* (isoform PA) was amplified by PCR form DGC clone RE12101 using WASp ORF primers (Table S2). The *comt* ORF was cloned from oligo dT primed cDNA using Comt ORF primers (Table S2). Both products were cloned into pCR8/GW/Topo (Invitrogen, CA), sequenced, and transferred into pTpaCW using Clonase L/R (Invitrogen) to give pTpaCW-WASp and pTpaCW-comt. Embryo injections were performed by Rainbow Transgenic Flies (Camarillo, CA, USA) and transgenics balanced using standard procedures.

For RT-qPCR analysis, fat bodies from wandering third-instar larvae of the appropriate genotypes were dissected in PBS, washed three times with PBS, and lysed in 0.8 mL TRIzol reagent (Invitrogen) according to the manufacturer’s instructions ^88^. Total RNA was isolated, and cDNA was synthesized using the SuperScript IV First-Strand Synthesis System (Thermo Fisher Scientific) according to the manufacturer’s instructions. Quantitative PCR was performed on a QuantStudio 6 Real-Time PCR System (Applied Biosystems) using SYBR Green PCR Master Mix (Thermo Fisher Scientific). Relative mRNA expression levels of *FAM21*, *Snx3* were determined by RT-qPCR and normalized to the housekeeping gene *RpL11* and *Gapdh2*. The average expression of these two housekeeping genes was used as the normalization factor for ΔCt calculations. Relative gene expression was calculated using the 2-ΔΔCt method as described by ^89^. The primers used for qPCR are listed in Table S2.

### e-MI assays, immunohistochemistry and reporter quantification

e-MI assays were performed as described ^22, 24^. Briefly, 8-10 virgin females were crossed with 10-12 males of appropriate genotypes on standard fly food at 25°C until late 2^nd^/early 3^rd^ instar. Larvae were washed 3 times in H_2_O, and the sensor photoactivated by exposure to 405-nm light for 10 min on ice in 800 µl Graces medium (Invitrogen, 11,605–094) containing 10% heat-inactivated FBS (Atlanta Biochemicals, S11050). After photoactivation larvae were washed 3 times in H_2_O and split to 35 mm cell culture dishes with 4 filter papers containing 1 ml of 20% sucrose only (starvation) and 20% sucrose with heat-inactivated yeast (fed). For the drug treatment, 20% sucrose with heat-inactivated yeast was supplemented with 20mM paraquat (Sigma, 856,177–1 G; dissolved in H_2_O) or 50 µM etoposide (Sigma E1383; dissolved in DMSO). Approximately 15-20 3^rd^ instar larvae were processed were inverted, fixed in freshly made, 4% paraformaldehyde (Fisher ICN15014601) in 1x phosphate-buffered saline (PBS; Corning Cellgro, 55–031-PC) at 4°C overnight. After 3 washes in 1x PBS, fat body lobes of approximately 10 larvae were mounted in 20 µl DAPI fluoromount-G (Southern Biotech 0100–20) per slide.

Immunohistological stainings were done as published ^22, 24^. Briefly, early third instar larvae were washed thoroughly in water, cut open, turned inside out and fixed in freshly made 4% paraformaldehyde in 1xPBS for 1 h at room temperature or overnight at 4°C. Antibodies: mouse anti-Rab7 (DSHB; 1:20), secondary antibody Alexa Fluor 647-conjugated anti-mouse (Life Technologies).

Reporter puncta were quantified from images taken under similar conditions using Fiji/ImageJ software (NIH). The SPARQ plugin was used to count puncta ^90^. Briefly, channels were split, thresholded for puncta, and puncta count and number of nuclei were recorded. For clonal analysis, regions of interest (ROIs) were outlined, and puncta were counted using Fiji/ImageJ, omitting puncta at the edges of the ROIs. The image was then split into channels with the same ROI applied, and puncta were counted as above. The results were expressed as mean values ± SEM. Statistical analyses (one-way ANOVA with Tukey’s or Dunnett’s post hoc tests) were performed using GraphPad Prism 11. “n =” indicates fields of view unless noted differently (because FB are bilaterally symmetrical it is a reasonable estimate that the number of animals is minimally about n/2 and maximally equal to n).

### Clonal Analysis

Flp-FRT technique was used to generate mutant clones lacking GFP by heat shocking larvae at 37 °C for 45 min within 8 h of egg laying as described ^22, 24^. Flip-on–based overexpression clones were generated using *hsFlp act>CD2>Gal4 UAS-GFP* was combined with *r4-KFERQ-PAmC* ^24^. Overexpression clones were generated without heat shock making use of the leaky expression of the *hsFlp* promoter. Larvae were dissected at the indicated stages and processed for imaging as above.

## Abbreviations

ALR: autolysosome reformation
Arp: actin related protein
Atg: Autophagy related
CCDC53: Coiled-coil domain-containing protein 53 (aka. WASHC3)
CMA: chaperone mediated autophagy
Comt/NSF1: Comatose/N-ethylmaleimide-Sensitive Factor 1
ESCRT: endosomal sorting complex required for transport
e-MI: endosomal microautophagy
Eto: etoposide
F: fed
FAM21: Family with sequence similarity 21 (aka. WASHC2; WASH complex subunit 2)
FB: Fat body
hps: hours post starvation
HSPA8/HSC70: heat shock cognate protein 70
LAMP: lysosome associated membrane protein
MA: macroautophagy
mC: mCherry
mTORC1/Tor: mechanistic target of rapamycin complex 1
NMJ: neuromuscular junction
NPF: actin nucleation promoting factor
PIP₂: phosphatidylinositol 4,5-bisphosphate
PQ: paraquat
ROS: reactive oxygen species
Snx: Syntaxin
St: starved
SWIP: Strumpellin and WASH-interacting protein (aka. WASHC5)
VCA: verprolin-homology/central/acidic domain
Vps: Vacuolar Protein Sorting-associated protein
WASH (washout): Wiskott-Aldrich syndrome protein and SCAR homolog (aka. WASHC1)
WASp: Wiskott Aldrige Syndrome protein
WHAMM/Whamy: WASP homolog-associated protein with actin, membranes, and microtubules.

## Data availability

This study includes no data that needs to be deposited in external repositories.

## Author contributions

SS and AJ designed the study, analyzed the data and wrote the manuscript. SS performed the experiments.

## Acknowledgements

We thank Drs. T. Neufeld (University of Minnesota), T. E. Rusten (Radium Hospital, Oslo, Norway), G. Juhasz (Szeged, Hungary), T. Vaccari (University of Milan, Italy), M. Mlodzik (Mount Sinai, NY), V. Tsarouhas (Stockholm University, Sweden), and the Bloomington Stock and Vienna *Drosophila* Resource Centers for kindly sharing fly strains, as well as Einstein Analytical Imaging Facility for microscope use supported by NCI grant P30CA013330 and SIG #1S10OD023591-01. We thank Dr. A. Petsakou for critically reading the manuscript. This work was supported by a research grant from the American Parkinson Disease Association (APDA 1058594 to S.S.) and NIH/NIGMS grant GM119160 (to A.J.).

## Figure legends

**Figures S1:** (**A**) Quantification of e-MI puncta per cell at 4hps under fed (F; grey) and starved (St.; magenta) conditions in indicated genotypes. At 4hps, compared to PAmC-WASp alone, knockdown of *Atg1* blocks the induction of early e-MI due to requirement of Atg1 for starvation induction. In contrast, knock-down of *Atg7* does not affect WASp induced e-MI. **(B)** Schematic representation of *Drosophila* NPF family proteins WASp, Wash and SCAR. Conserved domains are indicated as follows: WH, WASp homology domain; B, basic domain; GBD, GTPase-binding domain; P, proline-rich region; VCA, Verprolin, connecting and acidic domain responsible for Arp2/3 complex binding and activation. Numbers indicate length in amino acids. **(C, D)** The Arp2/3 complex is not required for autophagosome formation in larval FB . Cells mutant for *Arp2^A^* (lacking GFP expression and marked by a dotted yellow line) show comparable number of mCherry-Atg8a puncta to surrounding wildtype cells at 4hps (quantified in D). Monochromatic images show the indicated channels; scale bar: 20 µm. (A) One-way ANOVA (Tukey) p < 0.0001; (D): Student t-test; *** p< 0.001; **** p< 0.0001; n: number of fields of view in (A) and cells in (D).

**Figure S2:** Cdc42 is dispensable for e-MI. **(A-D)** Compared to fed conditions (A, B), 25h starvation induces e-MI in 3^rd^ instar larval fat body (C) that is not lost upon RNAi mediated knockdown of *Cdc42* (D). **(E)** Quantification of puncta number per cell of indicated genotypes at 25h under fed (gray) and starved (magenta) conditions. Monochromatic images show the indicated channels; nuclei are in blue; scale bar: 20 µm. One-way ANOVA (Tukey) p< 0.0001; *** p< 0.001; **** p< 0.0001; n: number of fields of view.

**Figure S3**: Regulation of e-MI by Wash is not restricted to starvation. Quantification of e-MI induction upon *cg>wash* expression at 4h of treatment. Feeding larvae with the DNA-damaging agent etoposide (Eto) or the oxidative stress inducer paraquat (PQ) induces robust e-MI puncta upon Wash overexpression. Student’s t-tests; * p< 0.05; *** p< 0.001; n: number of fields of view.

**Figure S4:** e-MI is independent of Rho-GTPases. **(A)** Expression of dominant-negative Rho1 *(UAS-Rho1^N^*^19^) was induced in flip-on clones in FBs expressing *r4-KFERQ-PAm*C. e-MI puncta formation (A’) was comparable between *Rho1^N^*^19^ cells (coexpressing GFP; A”) and surrounding wild type cells. Expressing cells are outlined by a dotted yellow line. **(B)** Cells mutant for *Rho1^M^*^1^ in mosaics (lacking GFP expression) show no change in e-MI sensor puncta formation at 25hps compared to neighboring wildtype cells. Mutant clones are outlined by a dotted yellow line. **(C)** Expression of dominant-negative Rac1 *(UAS-Rac1^N^*^17^) was induced in flip-on clones in FBs expressing *r4-KFERQ-PAm*C. At 25hps, sensor puncta formation was comparable between *Rac1^N^*^17^ expressing (marked by GFP and surrounded by a yellow dotted line; C”) and surrounding wild type cells. Monochromatic images show the indicated channels. Nuclei are in blue.

**Figure S5:** qPCR showed that *FAM21^IR^ ^GLC^*^01693^ and *Snx3^IR^ ^HMS^*^02344^ knockdown using *cg-Gal4* reduces the corresponding mRNA levels to 4.3% and 1.1% of WT level, respectively.

**Figure S6:** Expansion of the Rab7 compartment upon Wash overexpression. **(A-B)** Compared to control (A), Wash overexpression (B) results in the induction of expedited KFERQ-PAmC sensor puncta formation (B’’) as well as expansion of (late)endolysosomes marked by GFP-HsLAMP1 (B’) and of late endosomes stained for by anti-Rab7 (B”) at 4hps. Areas marked by yellow square in (A) and (B) are shown enlarged in (1) and (2), respectively. (C) Quantification of puncta number per cell at 4hs of indicated conditions and markers. C: control; W: *Wash* expression. Monochromatic images show the indicated channels. Nuclei are in blue. Scale bars: 20 µm (A, B) and 5 µm(1, 2). Student’s t-tests; * p< 0.05; ** p< 0.01; *** p< 0.001; ns: not significant; n: number of fields of view.

