## Supplemental Table 1 for "Actin nucleation promoting factors drive Arp2/3 dependent endosomal microautophagy"

**Supporting Table 1 Genotypes**

**Genotypes:**

**Figure 1**

**A, C, F:** $w^-$; cg-Gal4/+; UAS-PAmCherry-Comt#wt17/+

**B, D, G:** $w^-$; cg-Gal4/+; UAS-PAmCherry-WASp#wt49B/+

**E, H:** $w^-$; cg-Gal4/+; UAS-PAmCherryComt#wt17/+

$w^-$; cg-Gal4/+; UAS-PAmCherryWASp#wt49B/+

**I:** hsflp122/$w^-$ or Y; cg-Gal4 FRT42D UAS-GFPnls / FRT42 $Vps25^{A3}$; UAS-PAmCherryComt#17/+

**J:** hsflp122/$w^-$ or Y; cg-Gal4 FRT42D UAS-GFPnls / FRT42 $Vps25^{A3}$; UAS-PAmCherryWasp#49B/+

**K:** hsflp122/$w^-$ or Y; cg-Gal4 FRT42D UAS-GFPnls / FRT42 $Vps32^{G5}$; UAS-PAmCherryWasp#49B/+

**Figure 2**

**A, C, E (control):** $w^-$; cg-Gal4 UAS-GFP-HsLAMP1(RC-4)/+; UAS-KFERQ-PA-mCherry 3B/+

**B, D, E (WASp):** $w^-$; cg-Gal4 UAS-GFP-HsLAMP1(RC-4)/+; UAS-KFERQ-PA-mCherry 3B/ UAS-WASp

**F:** $w^-$; cg-Gal4/+; UAS-mCherry-Atg8a/+

$w^-$; cg-Gal4/+; UAS-mCherry-Atg8a/ UAS-WASp

**G:** hsflp122/$w^-$ or Y: cg-Gal4 FRT42D UAS-GFPnls / FRT42 $Vps32^{G5}$; UAS-PAmCherryWasp#49B/+

**H:** $w^-$; cg-Gal4/+; UAS-PAmCherry-WASp#wt49B/ +

**I:** $w^-$; cg-Gal4/+; UAS-PAmCherry-WASp#wt49B/ UAS-$Cdc42^{IR\ HMS01502}$

**J:** $w^-$; cg-Gal4/+; UAS-PAmCherry-WASp#wt49B/ +

$w^-$; cg-Gal4/+; UAS-PAmCherry-WASp#wt49B/ UAS-$Cdc42^{IR\ HMS01502}$

**Figure 3**

**A, B:** hsflp122/$w^-$ or Y; UAS-KFERQ-PAmCherry 19B/ +; r4-Gal4 FRT82B UAS-GFPnls /FRT82B $WASp^{1}$

**C:** $w^-$; cg-Gal4/+; UAS-KFERQ-PA-mCherry 3B/+

**D:** $w^-$; cg-Gal4/ SCAR- GFP-SCAR; UAS-KFERQ-PA-mCherry 3B/ +

**E:** UASp-GFP-Wash/ $w^-$; cg-Gal4/+; UAS-KFERQ-PA-mCherry 3B/ +

**F:** $w^-$; cg-Gal4/+; UAS-KFERQ-PA-mCherry 3B/+

UASp-GFP-Wash/ $w^-$; cg-Gal4; UAS-KFERQ-PA-mCherry 3B/ +

$w^-$; cg-Gal4/ SCAR- GFP-SCAR; UAS-KFERQ-PA-mCherry 3B/ +

**Figure 4**

**A:** $w^-$; cg-Gal4 UAS-GFP-HsLAMP1(RC-4)/+; UAS-KFERQ-PA-mCherry 3B/+

**B:** $w^-$; cg-Gal4 UAS-GFP-HsLAMP1(RC-4)/ $SCAR^{IR\ HMC03361}$; UAS-KFERQ-PA-mCherry 3B/+

**C:** $w^-$; cg-Gal4 UAS-GFP-HsLAMP1(RC-4)/ $wash^{IR\ HMC05339}$; UAS-KFERQ-PAmCherry 3B/+

**D:** $w^-$; cg-Gal4 UAS-GFP-HsLAMP1(RC-4)/+; UAS-KFERQ-PA-mCherry 3B/+

$w^-$; cg-Gal4 UAS-GFP-HsLAMP1(RC-4)/ $SCAR^{IR\ HMC03361}$; UAS-KFERQ-PA-mCherry 3B/+

$w^-$; cg-Gal4 UAS-GFP-HsLAMP1(RC-4)/ $wash^{IR\ HMC05339}$; UAS-KFERQ-PAmCherry 3B/+

**E:** $w^-$; cg-Gal4 UAS-GFP-HsLAMP1(RC-4)/+; UAS-KFERQ-PA-mCherry 3B/+

$w^-$; cg-Gal4 UAS-GFP-HsLAMP1(RC-4)/ $wash^{IR\ HMC05339}$; UAS-KFERQ-PAmCherry 3B/+

$w^-$; cg-Gal4 UAS-GFP-HsLAMP1(RC-4)/ +; UAS-KFERQ-PAmCherry 3B/ $wash^{IR\ GD024642}$

**Figure 5**

**A, E:** UASp-Arp3-GFP/ $w^-$; cg-Gal4/+; UAS-KFERQ-PAmCherry 3B/+

**B, F:** $Arp2^{A}$ FRT19A/ hsflp122 Ubi-GFP FRT19A; cg-Gal4/+; UAS-PAmCherry-WASp#49B/+

**C, G:** $Arp2^{A}$FRT19A/ hsflp122 Ubi-GFP FRT19A; cg-Gal4/ UAS-KFERQ-PAmCherry 19B; +/+

**D:** hsflp122/ $w^-$ or Y; $Arpc1^{Q25sd}$ FRT40A/ UAS-2xeGFP FRT40A FbGal4; UAS-KFERQ-PAmCherry 3B/+

**Figure 6**

**A:** $w^-$; cg-Gal4/+; UAS-KFERQ-PA-mCherry 3B/+

**B:** $w^-$; cg-Gal4/+; UAS-KFERQ-PAmCherry 3B/FAM21$^{IR\ GLC01693}$

**C:** $w^-$; cg-Gal4/+; UAS-KFERQ-PAmCherry/+

$w^-$; cg-Gal4/+; UAS-KFERQ-PAmCherry 3B/FAM21$^{IR\ GLC01693}$

$w^-$; cg-Gal4/+; UAS-KFERQ-PAmCherry 3B/Snx$^{IR\ HMS02344}$

**D:** hsflp122/ w- or Y: cg-Gal4 FRT42D UAS-GFPnls / FRT42D Vps35$^{MH20}$; UAS-KFERQ-PAmCherry 3B/+

**Supplementary Figures**

**Figure S1**

**A**: $w^-$; cg-Gal4/+; UAS-PAmCherry-WASp#wt49B/+

$w^-$; cg-Gal4/ UAS-Atg1$^{IR\ GD16133}$; UAS-PAmCherry-WASp#wt49B /+

$w^-$; cg-Gal4; +/ UAS-PAmCherry-WASp#wt49B /UAS-Atg7$^{IR\ GD45558}$

**C,D**: Arp2$^A$ FRT19A/ hsflp122 Ubi-GFP FRT19A; Atg8a>3xmCherryAtg8a/+; +/+

**Figure S2:**

**A, C, E**: $w^-$; cg-Gal4/+; UAS-KFERQ-PAmCherry/+

**B, D, E:** $w^-$; cg-Gal4/+; UAS-KFERQ-PAmCherry/ UAS-Cdc42$^{IR\ HMS01502}$

**Figure S3:**

$w^-$; cg-Gal4/+; UAS-KFERQ-PAmCherry 3B/+

$w^-$; cg-Gal4/ UAS-wash; UAS-KFERQ-PAmCherry 3B/ +

**Figure S4:**

**A:** hsflp122 act>CD2>Gal4 UAS-GFP / w- or Y; r4-KFERQ-PA-mCherry 8/ UAS-Rho1$^{N194.3}$

**B:** hsflp122/+; cg-Gal4 FRT42D UAS-GFPnls / FRT42 Rho1$^{M1}$; UAS-KFERQ-PA-mCherry 3B

**C:** hsflp122 act>CD2>Gal4 UAS-GFP / w- or Y; r4> KFERQ-PA-mCherry 8 /UAS-Rac1 $^{N17}$

**Figure S5:**

$w^-$; cg-Gal4/+; UAS-KFERQ-PAmCherry 3B/+

$w^-$; cg-Gal4/+; UAS-KFERQ-PAmCherry 3B/ FAM2 $^{IR\ GLC01693}$

$w^-$; cg-Gal4/+; UAS-KFERQ-PAmCherry 3B/ Snx3 $^{R\ HMS02344}$

**Figure S6:**

**A**: $w^-$; cg-Gal4/+: UAS-GFP-HsLAMP1/+; UAS-KFERQ-PAmCherry 3B/+

**B**: $w^-$; cg-Gal4/+: UAS-GFP-HsLAMP1/ UAS-wash; UAS-KFERQ-PAmCherry 3B/ +

**C**: $w^-$; cg-Gal4/+: UAS-GFP-HsLAMP1/+; UAS-KFERQ-PAmCherry 3B/+

$w^-$; cg-Gal4/+: UAS-GFP-HsLAMP1/ UAS-wash; UAS-KFERQ-PAmCherry 3B/ +
