## Supplemental Table 2 for "Actin nucleation promoting factors drive Arp2/3 dependent endosomal microautophagy"

Table S1. Primer sequences used in this study.

| Gene | Forward Primer (5′→3′) | Reverse Primer (5′→3′) |
|---|---|---|
| RpL11 | CTTACAAGTTTCCGCGAGGG | CGCAGATGTTCAGGCAGAGT |
| Gapdh2 | CTACCTGTTCAAGTTCGATTCGAC | AGTGGACTCCACGATGTATTCG |
| Snx3 | CGG GAT AGC AAG ATT GTG GT | GCT TTC GTC AAA GAT GCC CT |
| Fam21 | GAT GAA GCG CAT TTC CCA GAA TC | CCA CCC GAT ACT CCA CGA AC |
| PAmCherry | TTGTATACCGGTGCTTGTACAGCTCGT CCATGCC | TACACCTAGGCGGTACCACTGCAGTGAAT TCGGAGCTCCGCCACCATGGTGAGCAAG GGCGAGGAG |
| Comt ORF | ATGGCTTATATTTTGAAGGCCA | TTACTGCCGCGCCACCATGTC |
| WASp ORF | ATGAGCAGCGGAATGAGGTCG | TTAGTCCCACTCCCCTTCGTT |
| Wasp1 rec | AAGGGATTTGATCTGGCGGG | AGCATGTTTAAAATGGCTAAGCG |
