## Supplementary figures and images for "Actin nucleation promoting factors drive Arp2/3 dependent endosomal microautophagy"

### Supplemental Figure 1

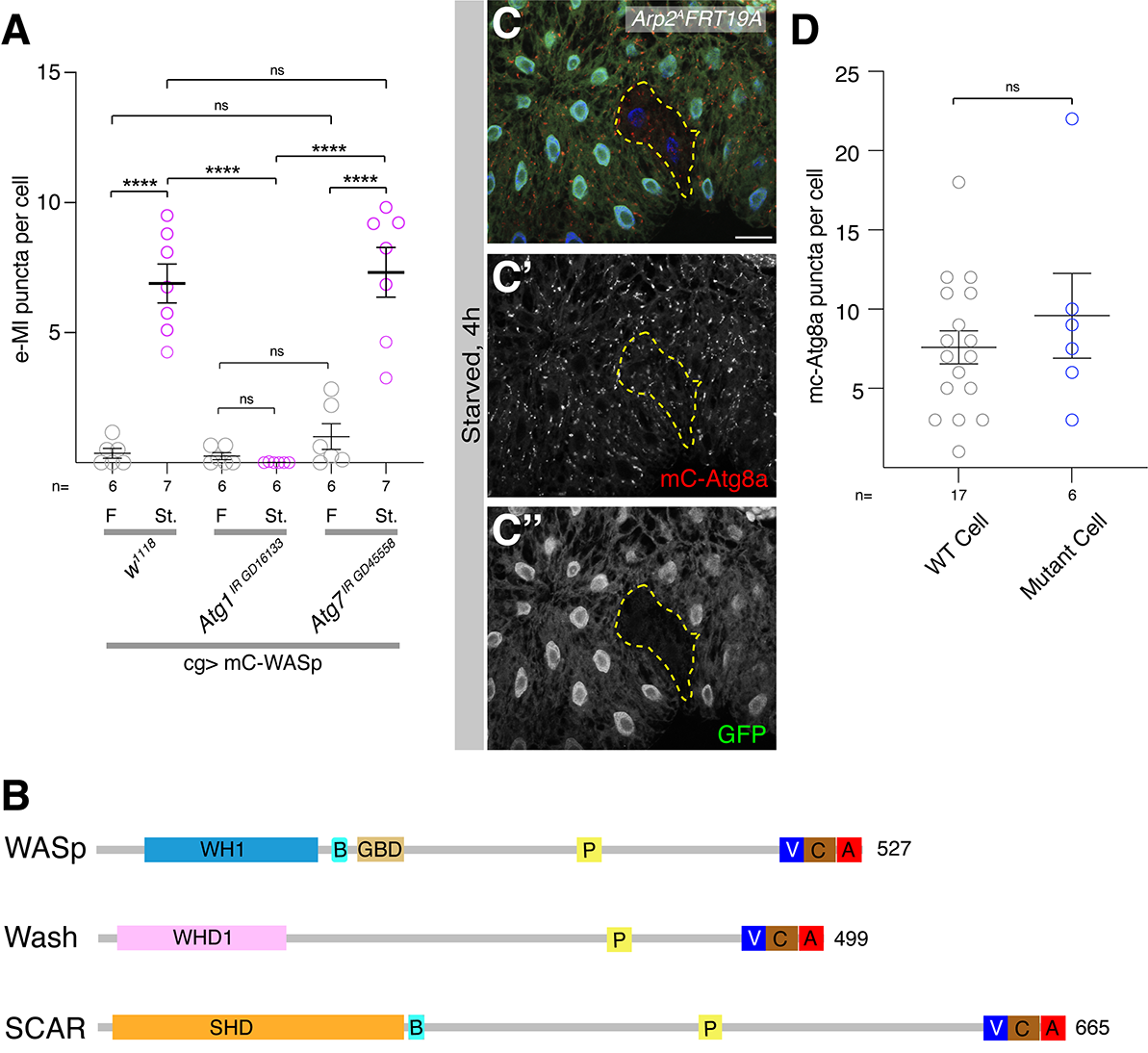

### Supplemental Figure 2

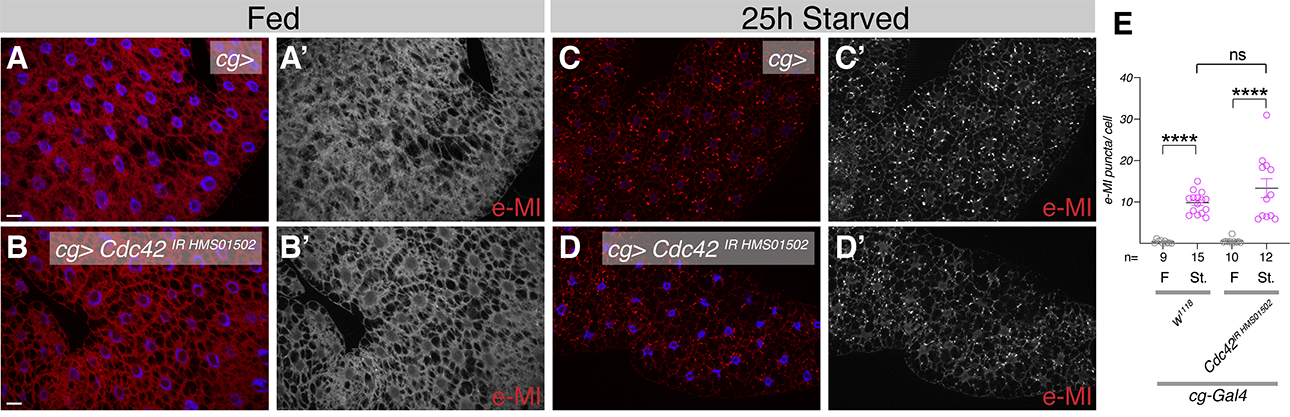

### Supplemental Figure 3

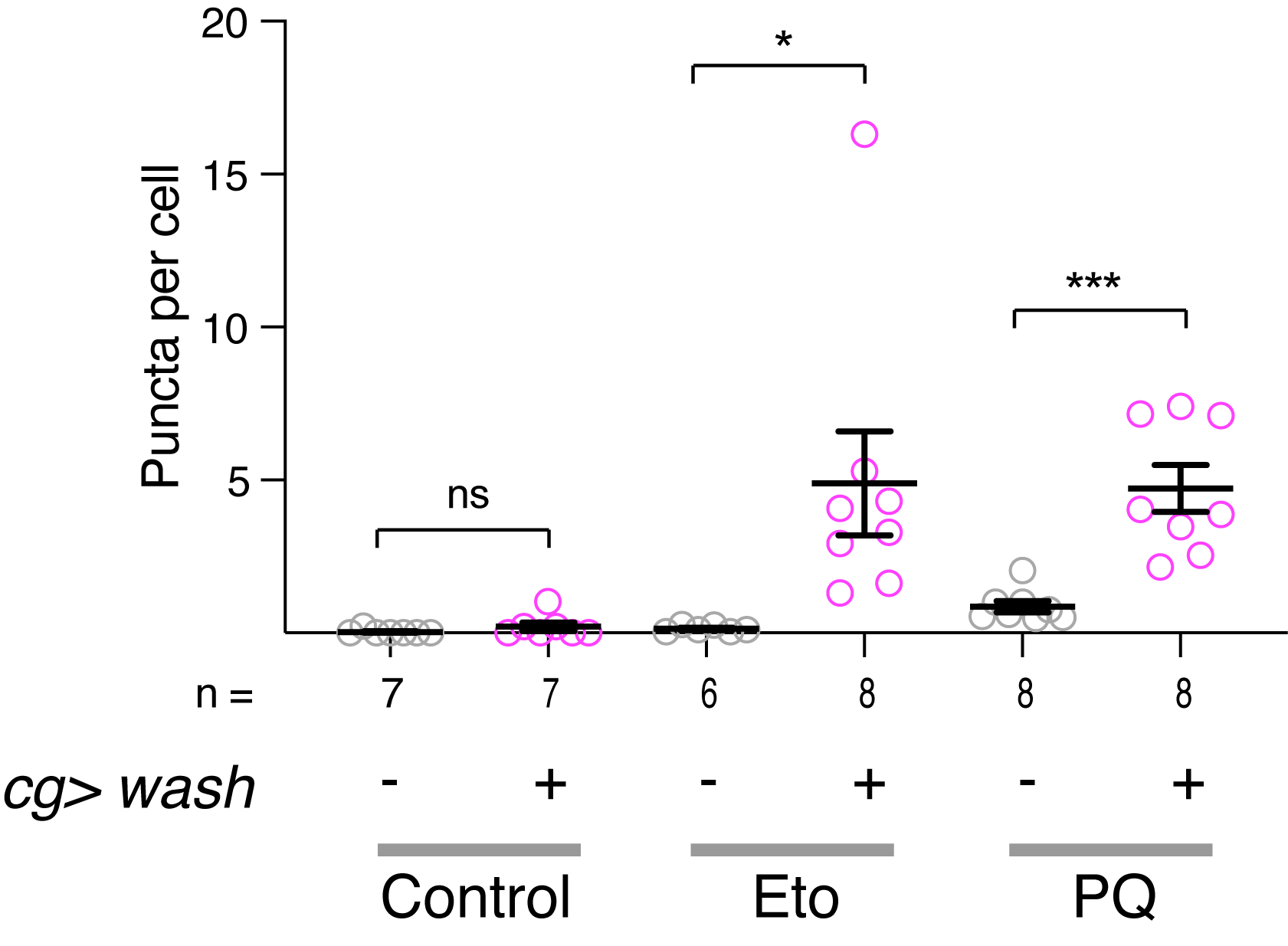

### Supplemental Figure 4

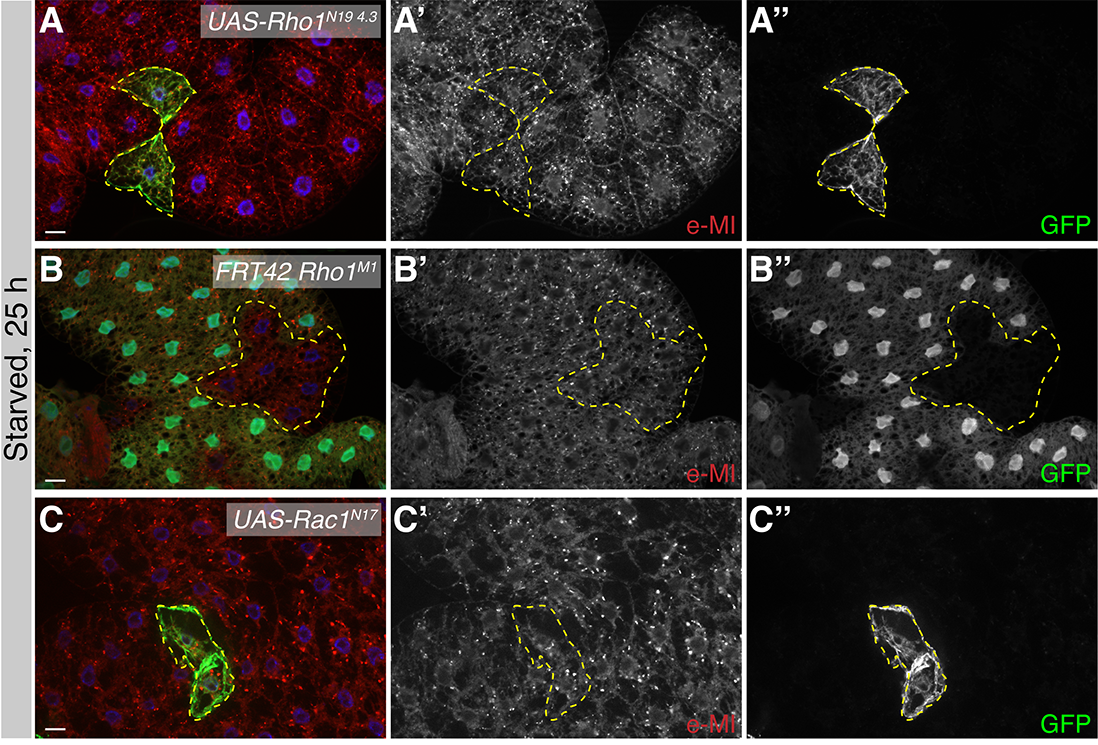

### Supplemental Figure 5

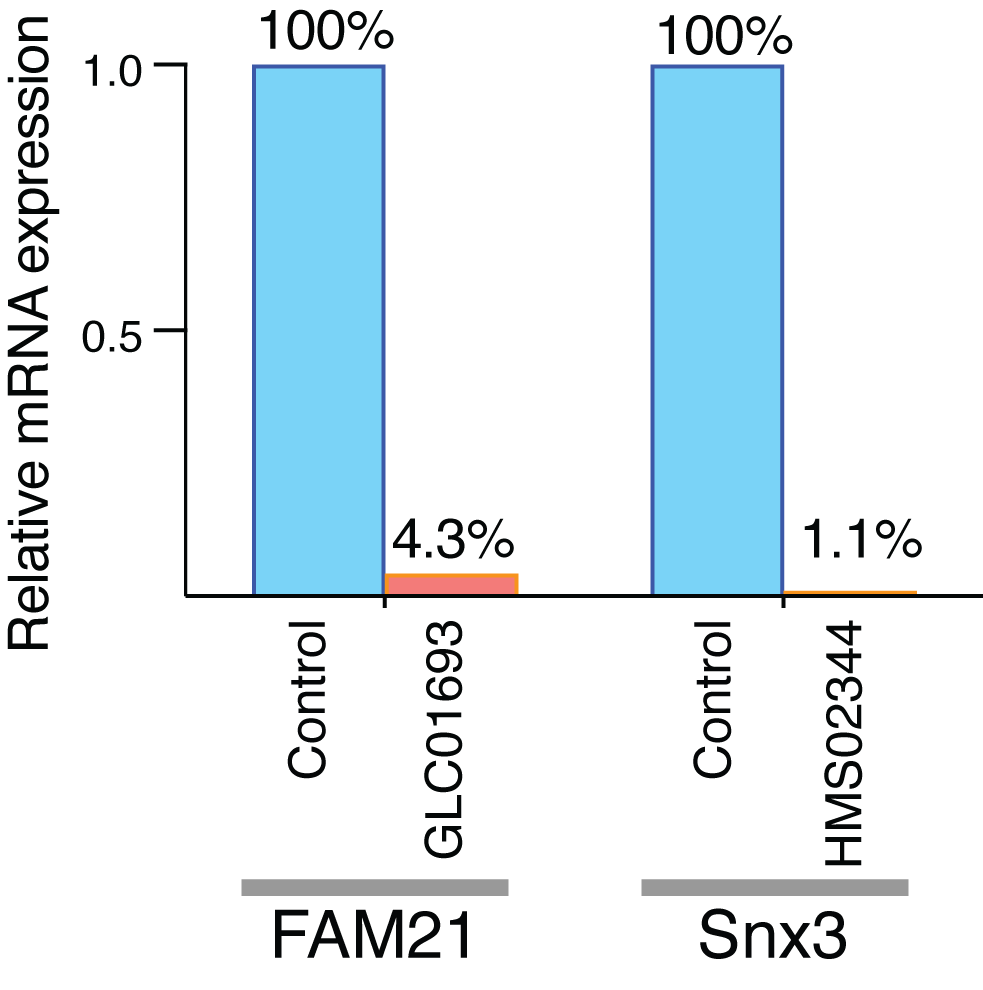

### Supplemental Figure 6

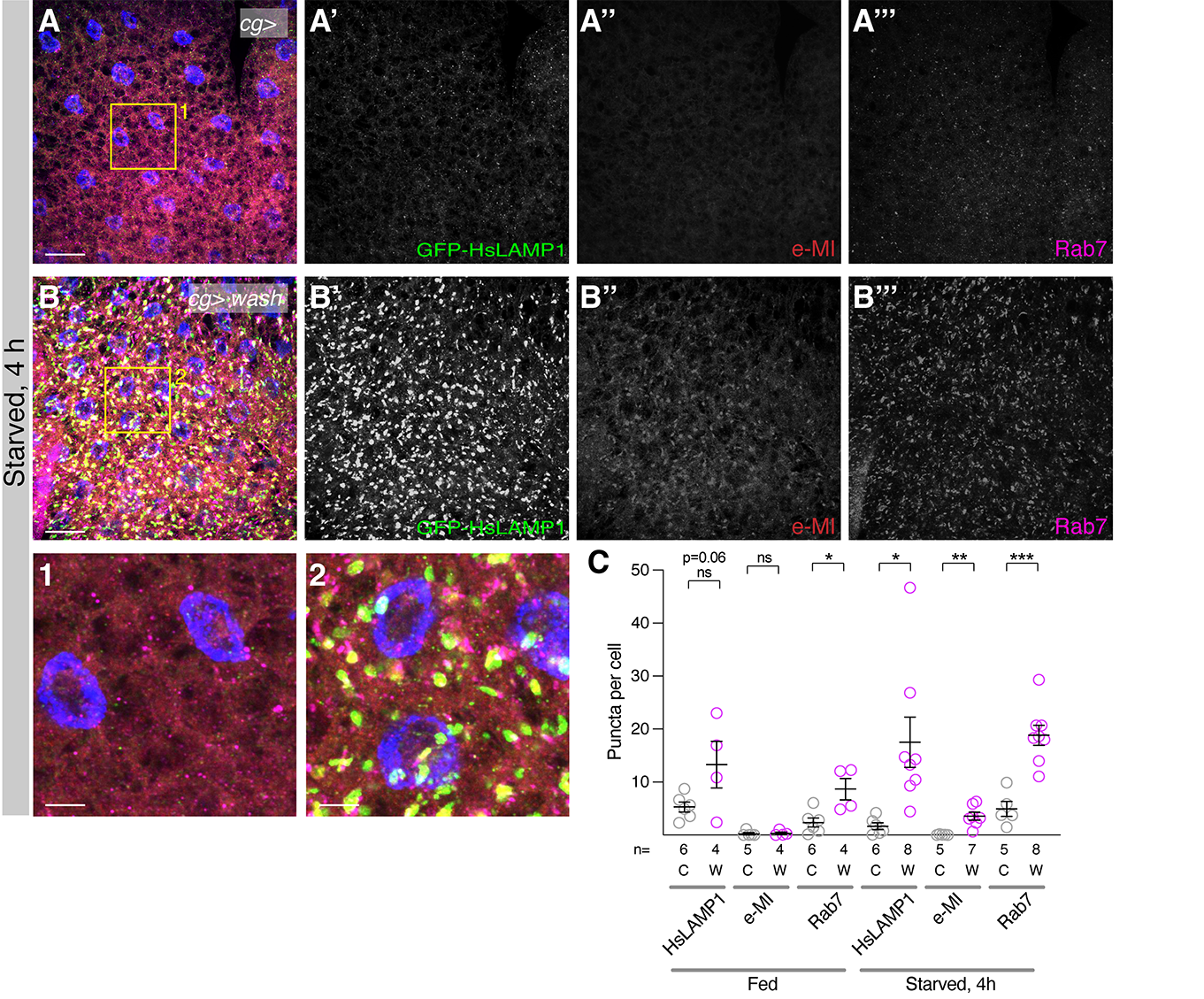
